# Fast Grip Force Adaptation To Friction Relies On Localized Fingerpad Strains

**DOI:** 10.1101/2021.07.20.452911

**Authors:** Félicien Schiltz, Benoit P. Delhaye, Frédéric Crevecoeur, Jean-Louis Thonnard, Philippe Lefèvre

## Abstract

Humans can quickly adjust their grip force to a change in friction at the object-skin interface during dexterous manipulation in a precision grip. To perform this adjustment, they rely on the feedback of the mechanoreceptive afferents innervating the fingertip skin. Because these tactile afferents encode information related to skin deformation, the nature of the feedback signaling a change in friction must somehow originate from a difference in the way the skin deforms when manipulating objects of different frictions. To better characterize the origin of the underlying sensory events, we asked human participants to perform a grip-lifting task with a manipulandum equipped with an optical imaging system. This system enabled to monitor fingertip skin strains through transparent plates of glass that had different levels of friction. We observed that, following an unexpected change in friction, participants adapted their grip force within 370ms after contact with the surface. By comparing the deformation patterns when unexpectedly switching from a high to a low friction condition, we found a significant increase in skin deformation inside the contact area arising over 100ms before the motor response, during the loading phase, suggesting that local and partial deformation patterns prior to lift-off are used in the nervous system to adjust the grip force as a function of the frictional condition.

## Introduction

In a seminal paper in the 80s, Johansson and Westling described how efficiently human participants handle objects of different textures and friction (Johansson and Westling, 1984). They observed that when lifting objects, humans scaled their grip force (GF) to the frictional properties of the surface, such that an object with a slippery surface was gripped more firmly than one with a sticky surface. Moreover, it was found that a change in the frictional properties of the object from one trial to the next elicited a GF adaptation that was observable only 100ms after contact with the surface. Such adaptation was cancelled under local anesthesia, underlining the essential role of afferent feedback (Westling and Johansson, 1987; Nowak et al., 2001; Augurelle et al., 2003; Witney et al., 2004). A rapid feedback loop is thus able to take into account tactile afferent information about the surface efficiently (Johansson and Flanagan, 2009; Delhaye et al., 2018).

However, the mechanisms underlying such a feedback loop remain unknown. Indeed, since the surfaces used in the aforementioned paper (Johansson and Westling, 1984) had very different textures, it is not clear whether the feedback provided by the afferents was related to the topography of the material, thereby quickly eliciting the recall of a motor memory related to the surface, or if the feedback was directly related to friction, such that the motor system could scale the GF accordingly. Notably, it was later demonstrated that human are able to adapt to changes in friction (Birznieks et al., 1998), even those that are not directly associated with a change in texture ( Cadoret and Smith, 1996). In this study, different textures and coatings were used to show that humans adjust the level of GF to the coefficient of friction and not to the texture when lifting objects. These results suggest that the skin deformation during each lifting movement can be used to scale the grip force without necessarily requiring a full slip event.

The possibility that humans adjust their grip force quickly based on an estimate of friction is supported by recent imaging studies of fingertip deformation during loading. We and others have shown that localized partial slips take place at the object-finger interface during tangential loading (Levesque, 2002; Tada et al., 2006; André et al., 2011; Delhaye et al., 2014). Partial slips are associated with substantial skin deformation in the contact area (Delhaye et al., 2016) that trigger a strong afferent responses (Delhaye et al., 2021a), and may therefore signal an impending slip. Importantly, reducing friction accelerates the progress of partial slips and leads to an earlier discharge of the tactile afferents, which can potentially inform the central nervous system about the upcoming contact instability (Khamis et al., 2014; Delhaye et al., 2021a). Moreover, the perception of tactile slip seems to be induced by skin deformations associated to partial slip, since it is perceived before full slippage and is impeded when the amount of strains is diminished by applying a coating that reduces friction (Barrea et al., 2018). Furthermore, generating artificial skin strains at the contact interface with the object during lifting also lead to an increase in GF (Farajian et al., 2020).

Taken together, the aforementioned findings lead us to hypothesize that partial slip, or the associated skin deformations, are a sufficient sensory signal to adjust the GF to the friction condition during active manipulation. To test this hypothesis, we sought to describe and quantify where and when skin deformation associated to partial slip take place following an unexpected change of friction, and if those allow participants to adapt their GF to a change in friction that is not associated with a change in texture. To this end, we asked human participants to repeatedly grip and lift a manipulandum, while the friction was changed unbeknownst to the participant (Fig 1A-B). We found that participants adjusted the GF to a change in friction only 114ms after liftoff (370ms after contact was made with the surface), suggesting that most of the sensory information about the friction change was available before the object started moving. To further understand the mechanisms underlying such adjustments, we imaged, at the same time, the contact area between the index finger and the object. We used those images to track the skin strains resulting from partial slip (Fig 1F-G), and reveal a localized strain contrast after friction changes very early in the trial, i.e. before liftoff. Our findings thus support the hypothesis that humans make use of localized strain patterns to adjust the GF to unexpected changes in friction.

**Figure 1:**
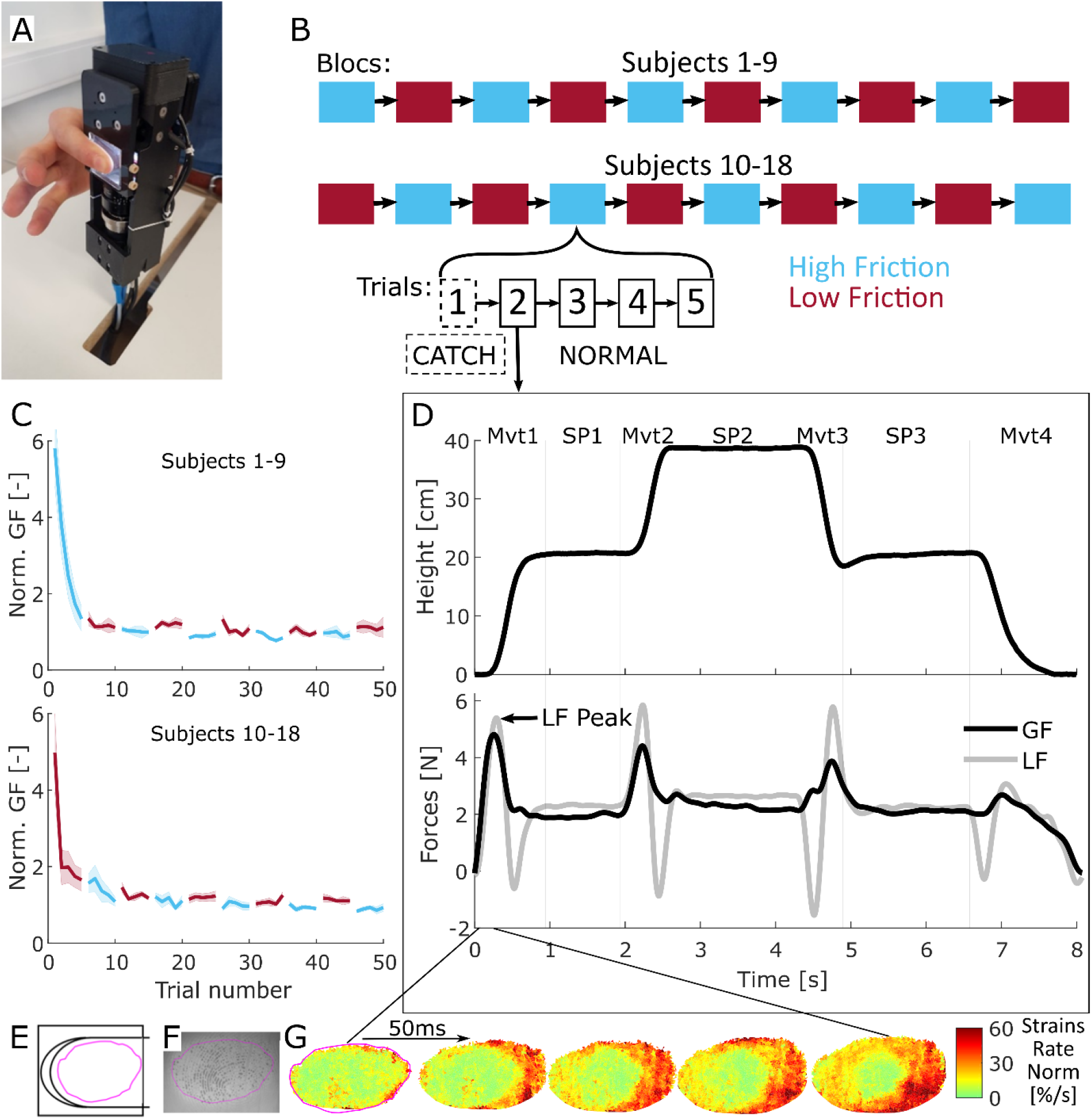
Experimental setup, experimental procedure, and typical trial. **A|** Participants held the manipulandum in a precision grip with both fingers applied on transparent glass plates. The device includes sensors allowing the measurement of the forces applied by both fingers as well as an imaging system allowing the recording of the index fingertip skin. **B|** Participants performed ten blocks of five trials. Transparent plates with high and low friction properties were interchanged between each block. Half of the participants started with the high friction material. The first trial of each block is called “CATCH” trial, as opposed to “NORMAL” trials. **C|** Group mean of the GF of the first SP for participants who started with the high friction material for the top graph and for participants who started with the low friction material for the bottom one. GF is normalized according to the procedure described in *Methods*. The shaded areas show the standard error of the mean. **D|** Evolution of the vertical position of the manipulandum and the forces applied during a typical trial. It consists of four fast movements (**Mvt**) with static phases (**SP**) in-between. Note that the second LF peak of each movement is due to the participant having to slow down the manipulandum because of the inertia of the system. The LF during the second static phase is slightly higher due to a larger portion of the cable being positioned under the device at this height. **E|** Position of the index finger and area of contact with the glass. All strains in this study are displayed as if they were observed through the glass during the manipulation. The distal side is on the left. The pink curve delimits the contact area. **F|**Typical image. Only the index finger was monitored. **G|** Heat maps of the norm of the skin strain rates obtained from a pair of consecutive images, as described in (Delhaye et al. 2016). This sequence shows deformations during the first 200ms of a typical first movement (Mvt1 in panel C). Strains are observed at the periphery of the contact area. The central stuck zone remains undeformed.

## Results

Participants performed a series of grip and lift trials using a custom-made manipulandum held in a precision grip (Fig 1A). Following an auditory cue, they were instructed to grip and lift the object vertically to reach a target (movement, “Mvt1” in Fig 1D), and then hold it stationary for one second (static phase, “SP1”). This task was followed by three point-to-point movements (Mvt2-4, and SP2-3, also paced by auditory cues, not analyzed here). After each experimental block consisting of five trials, the participants were asked to sit on a chair with their back turned away to the experimental set-up, and the experimenter quickly interchanged the surfaces without the participants noticing (Fig 1B). Two sets of glass plates having different levels of friction were used (see *Methods*). We defined the first trial following a surface change as a “catch trial”, since it included an unexpected change in friction, and the other four trials were called “normal trials”. The first two blocks were considered to be “training blocks” as the participants’ GFs decreased significantly during those for all participants and were thus excluded from the data analysis (Fig 1C).

The manipulandum was equipped with force sensors that allowed us to monitor the GF and the load force (LF, acting vertically and due to the object weight and inertia). A typical point-to-point movement was accompanied by two LF peaks, related to the acceleration and deceleration phases of the movement (Fig 1D). Note that each upward LF peak was paired with a GF peak. We synchronized all trials at the instant of the first LF peak. During the static phases, LF remained fairly constant (2.2N, the object’s weight) as did GF.

The index fingertip contact with the object’s surface was monitored through the glass plates using a high-speed, high-resolution camera (Fig 1E-F). Image processing techniques allowed us to track fingerprint movements and evaluate surface skin strains during the lift movement (Fig 1G, (Delhaye et al., 2014, 2016), see *Methods*). The rate of change of skin strains resulting from LF increase during the lift of the manipulandum were observed at the periphery of the contact area (Fig 1G). In most trials, the center of the fingertip remained stuck and non-deformed, except when full slip was reached (87 times out of 633, or equivalently 14% of trials). The contact area was an approximately ellipsoidal shape and increased over time following a logarithmic increase consistent with previous observations (André et al., 2011). On average, the contact area reached 85% of its maximum value 200ms after the time of contact and had usually reached its maximum value 340ms after initial contact (value reached: 94.4±5% of the maximum value, mean±std). This time corresponded to an average of 30ms prior to liftoff.

### Consistent difference in friction between smooth transparent materials

First, we verified that the two materials showed a consistent difference in their coefficient of friction. To that end, the friction between the fingertips, index and thumb, and the two sets of plates was measured at the end of the experiment to reduce potential cues about the different materials that could have been used during the manipulation task. Note that all participants except three reported that they did not notice that different materials were used. Following ((Barrea et al., 2016), see *Methods*), we characterized the coefficient of friction over a range of GFs relevant to our experiment (see Fig 2). The data were obtained for both fingers of all participants (and are reported in Supp Fig 1 for the index and Supp Fig 2 for the thumb) and fit with a negative power law. We observed that the coefficient of friction of the low friction glass remained lower than the one of the high friction glass across all levels of normal force tested, as shown in Fig 2A for a typical participant. From the fits obtained for each participant, we summarized the coefficient of friction by a single value, being the average coefficient of friction over the range of 1 to 5N (which spans the levels of GF used in this study, see e. g. Fig 1D), and across fingers. Overall, the friction was always higher in the case of the high friction material and the average relative difference was larger than 20% (paired t-test, t(17)=10.0696, p<0.001) (Fig 2B-C). Given that we aimed to observe behavioral adaption to changes in friction, we required a sufficient difference between materials and set a lower bound to a relative difference of 10%. Accordingly, one participant was removed from all subsequent analyses because the relative friction difference was too low (only 2%, less than the 10% required, see the unfilled dot in Fig 2B-C, participant S1 from Supp Fig 1–2). In summary, then, the two flat and transparent materials used in this study showed a consistent difference in friction, on average 23% across participants.

**Figure 2:**
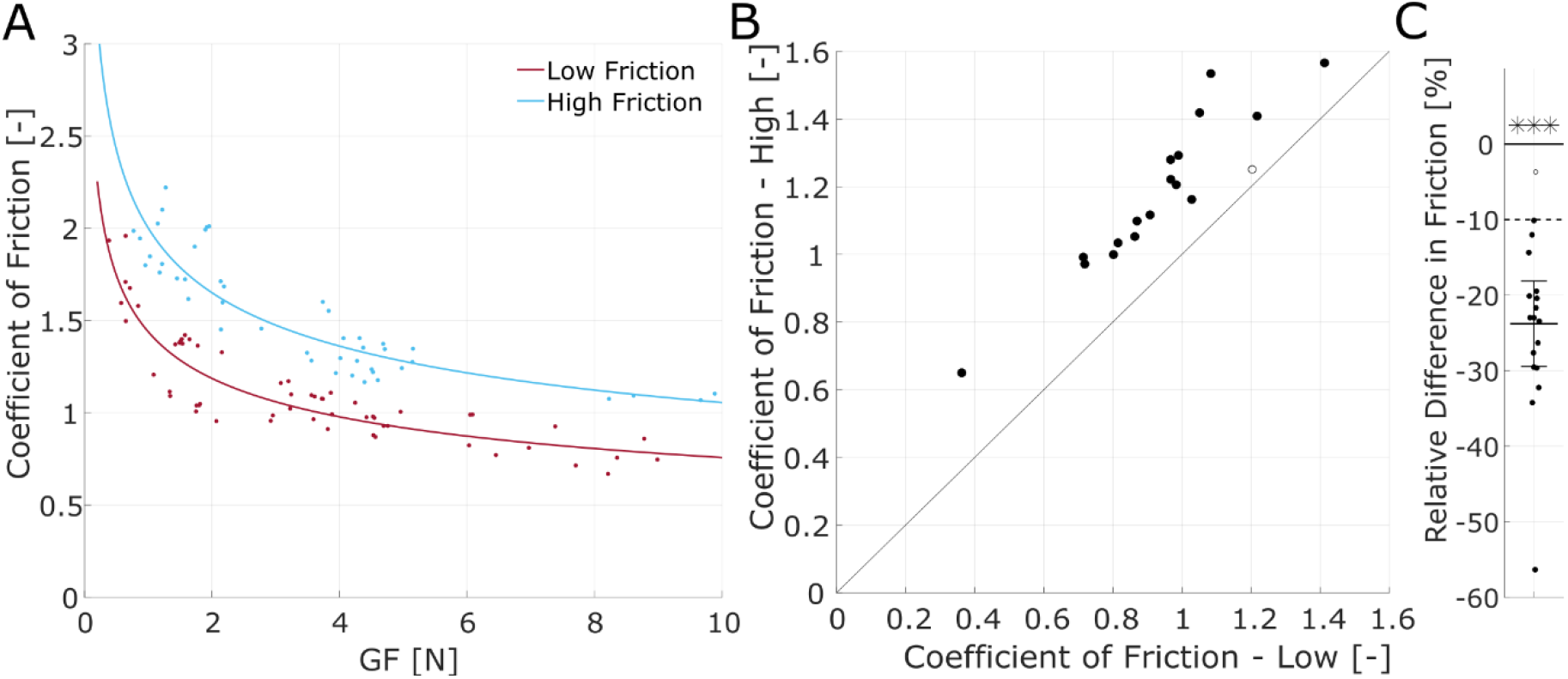
Consistent difference of friction. **A|** Coefficient of friction of the index finger as a function of grip force for a typical participant obtained using the method described in (Barrea et al. 2016). **B|** Mean coefficient of friction of both materials over the observed manipulation range (1-5N) for each participant (n=18). **C|** Difference in friction of the materials over the manipulation interval (1-5N), relative to the mean of the values of the coefficient of friction of both materials. The brackets show the 95% confidence interval of the mean. The dashed line indicates the level of sufficient difference in friction between materials for a participant to be included in the study. In panels B| and C|, one point corresponds to one individual participant and the data were averaged across both fingers and the empty circle shows the participant that was removed from the following analysis because of a difference of friction smaller than 10%.

### The grip force is adjusted to friction

Next, we sought to assess whether the different materials elicited different gripping behavior. First, we tested if participants could adapt to the difference in friction by using a consistently higher GF for the lower friction during the normal trials, i.e. those not following an unexpected change in friction. We found that indeed, most of the participants used a higher level of GF for the lower friction as averaged over the three static phases (Fig 3A), even though the level of GF varied widely across participants. Overall, the relative difference was statistically significant and close to 15 % (Fig 3B, mean 14.87±12.7%, paired t-test, t(16)=4.8224, p<0.001). Thus, participants spontaneously adjusted the GF level to the friction condition. Moreover, the relative GF difference was of the same order of magnitude as the relative friction difference. We observed that the participant that was removed from the analysis because of a lack of difference in friction showed no significant difference in behavior across the friction conditions (diff=2%, as measured by the average level of GF during the static phases of normal trials).

**Figure 3:**
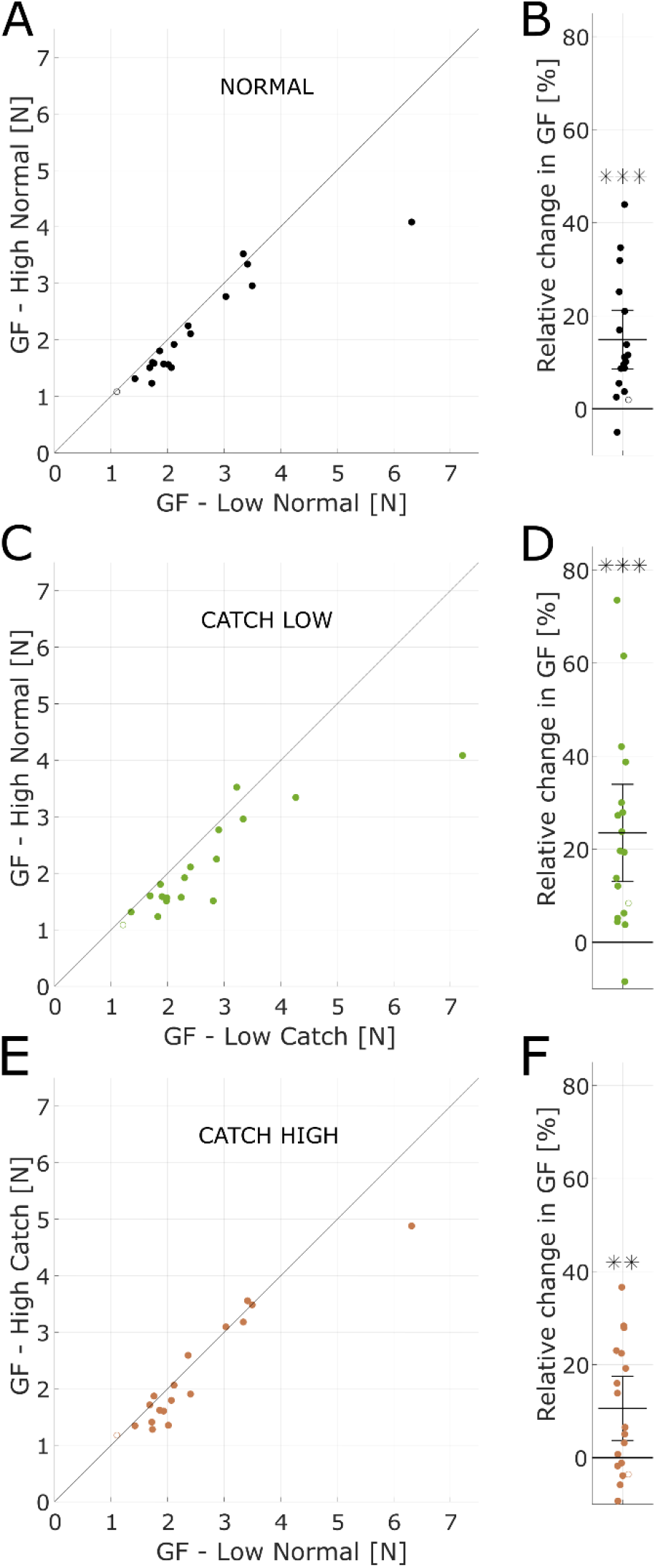
GF adaptation to friction during the static phases. **A|** Mean value of the GF for each material during the static phases of normal trials for all participants (n=18). **B|** Change in the mean value of the GF from the high to low friction material during the static phases of normal trials, relative to the mean value of the GF in both friction conditions. The brackets show the 95% confidence interval of the mean. **C, D|** Same as A, B|, except the static phases of the high friction normal trials are compared to the first static phase of the low friction catch trials, which directly follow. The bar delimited with hashed lines indicates the catch trials. **E, F|** Same as C, D|, except the static phases of the low friction normal trials are compared to the first static phases of the high friction catch trials, which directly follow. For D| and F|, the percentage of change is relative to the mean of the values of normal and catch trials. The participant that is removed from the study due to a low difference of friction is shown with an empty circle and is not included in the means and confidence intervals shown.

### Catch trials show GF adjustments within the first movement

Having observed that there was indeed a GF adaptation to friction for normal trials, we then tested if a change in friction elicited a quick adjustment of the GF, as observed in the first static phase of the catch trials. To that end, we compared the static GF after the first movement (SP1, Fig 1D) of the catch trials to the static GF experienced and learned during the normal trials of the preceding block of trials associated with the material with different frictional properties. The catch trials could be of two types: (1) low friction catch trials, following an adapted exposure to high friction, were referred to as “catch-low”, and (2) high friction catch trials, following an adapted exposure to low friction, were referred to as “catch-high”.

For the “catch-low” trials, which required an urgent increase in GF because the drop of friction increased the risk of slip and drop of the object, we found that the GF was already higher during the first static phase (Fig 3C-D, paired t-test, t(16)=4.5486,p<0.001)). The adaptation was close to 20%, thus already of the same magnitude as the adaptation learned throughout many trials (Fig 3B). Then, we looked at the “catch-high” trials, for which the urgency was lower since the experienced surface friction increased and therefore the risk of slippage was lower than for the directly preceding trials. We also found that the GF changed after only one movement, although the relative difference of GF was about 10% on average, thus not as large as for the catch trials in the other direction. Although this difference was not observed for all participants, it was statistically significant (Fig 3E-F, paired t-test, t(16)=3.1527,p<0.01).

In brief, participants adapted the GF to the friction condition, and this adaptation was already present at the end of the first movement following a catch trials, and for both directions of the changes in friction.

### Grip force adjustments start close to the time of liftoff of catch trials

Since we observed that the GF was already adjusted to the friction level after the first movement of catch trials, we investigated the temporal evolution of the GF during these catch trials to determine when the changes in GF evoked by the change in frictional properties arise (Figure 4). The participants produced movements with very similar kinematics across conditions, as shown by the height and LF curves, which could not be distinguished across conditions (Fig 4, top two rows of panels A and B). We observed that GF curves of the catch trials progressively diverged from those of the normal trials (Fig 4A-B, third row). In these graphs, the GF were normalized according to their average values during the first static phase for each subject (values reported in Supp Fig 3).

**Figure 4:**
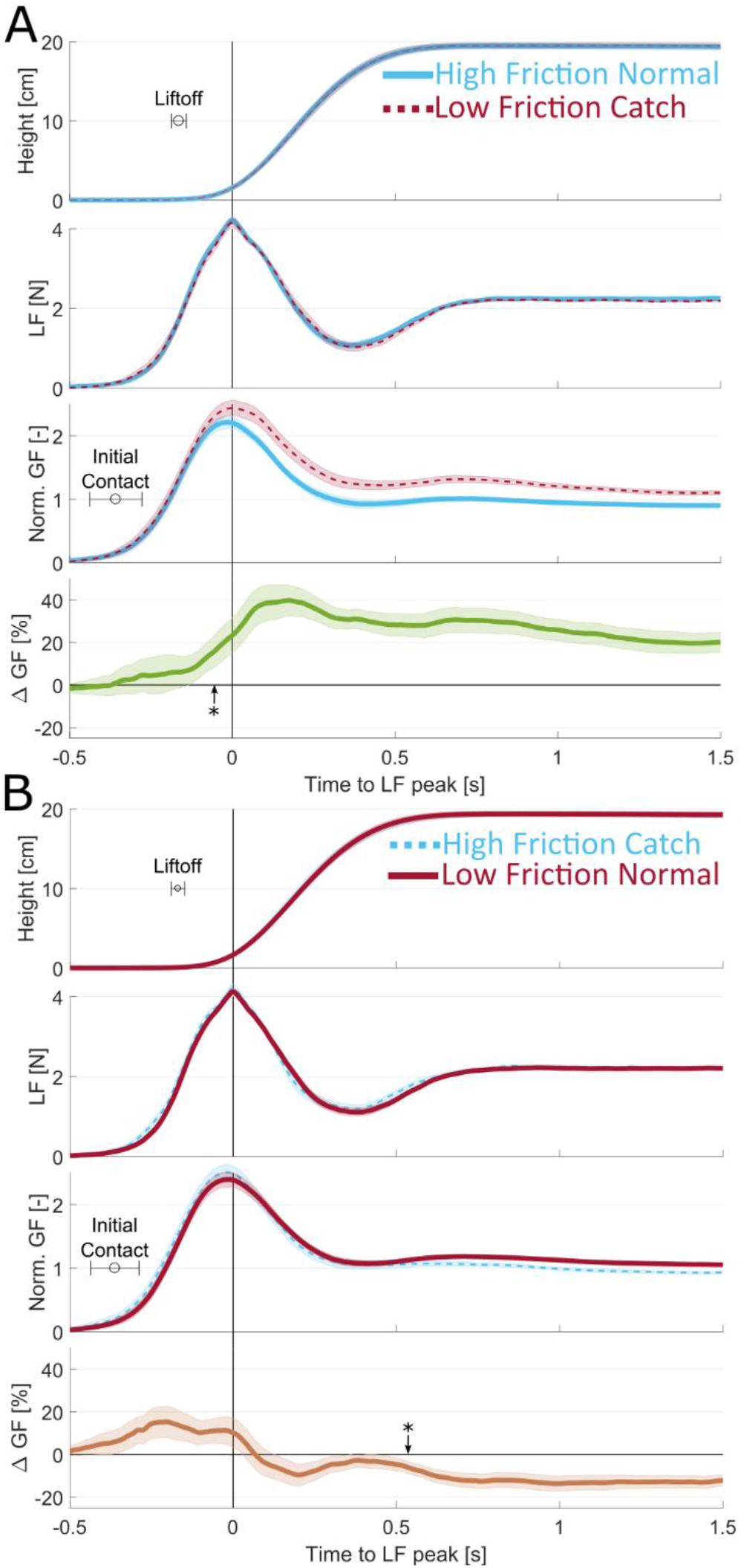
Adaptation to friction during the first movement of catch trials. **A|** Evolution of object height, LF, GF, and GF difference as a function of time for the first movement of catch-down trials for all participants (n=17). Trials are synchronized to the LF peak. Lines are averages across participants and shaded areas are the standard error of the mean. 0s indicates the time of the maximum of Load Force. Blue is for high friction and red is for low friction. Thick continuous traces are normal trials and dashed lines are catch trials. GF normalization values are reported in Supp Fig 3. The star shows the time of the statistically significant difference between GF curves (p<0.05). The lower panel shows catch minus normal trials. **B|** Same as A for the catch-down trials.

For the “catch-low” trials (Fig 4A), we found that the GF difference reached statistical significance very early, just after liftoff (114±23ms delay), or 50ms before the peak of LF (Fig 4A, bottom row). This timing corresponded to 308±80ms after initial contact. This difference was substantial, as it peaked on average at 40% with respect to GF during the first static phases of the normal trials.

For the “catch-high” trials (Fig 4B), the difference in GF reaches statistical significance later in the movement (540ms after the peak of LF, 709±20ms after liftoff, and about 902±74ms after initial contact) and is relatively smaller at the end of the first movement, as seen previously (Fig 3). Note that participants tended to apply a slightly higher level of GF at the very beginning of the contact of catch trials no matter the sign of the friction change, which might be explained by the short break between blocks. This systemic increase of the GF at the start of new blocks regardless of the actual friction condition suggests that participants were not able to anticipate the friction for the catch trials.

In summary, we showed that GF reaches a level that is significantly different from the level of the normal trials during the first lift following a change in friction, around the time of liftoff for the “catch-low” and a bit more than half a second later for the “catch-high” trials.

### Subtle contrast in skin strain rate before liftoff are cues for GF adjustments

Looking at the Δ GF traces (Figure 4A-B, bottom rows), we can observe an upward inflection for “catch-low” trials, and a downward inflection for “catch-high” trials, happening around the LF peak and suggesting that the corrective behavior kicks in around that time. To probe the timing of the reaction of the participants to the change in friction, we thus looked at the rate of change in GF (see Fig 5A).

**Figure 5:**
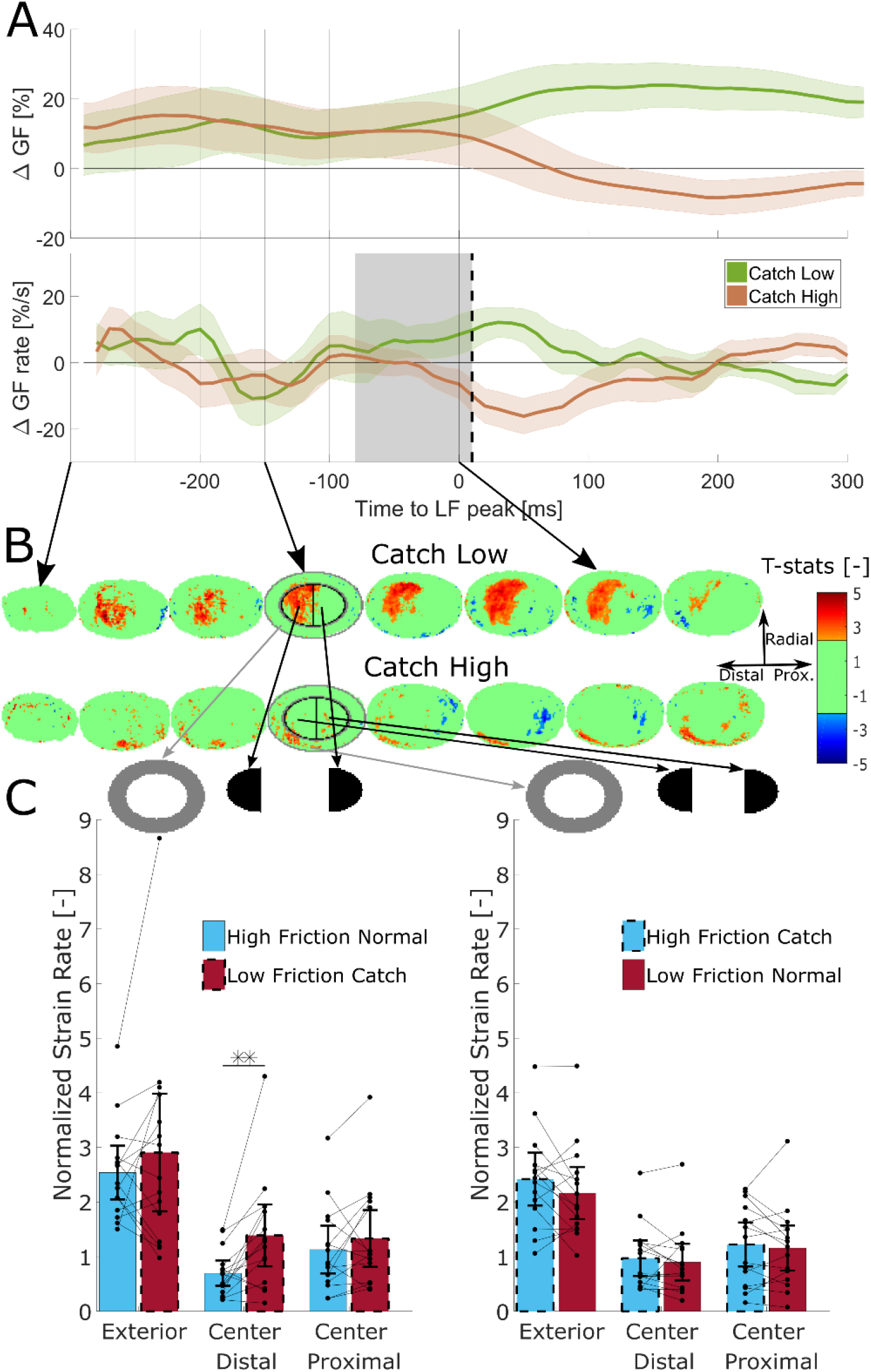
Differences in skin strains preceding the force adaptation. **A|** Mean difference of GF and mean difference of GF rate between catch and normal trials. Only participants for whom the images were of sufficient quality are included (n=15). 0ms indicates the time of the maximum of Load Force. Shaded areas are the standard error of the mean. The vertical dashed line indicates the time when the differences of Grip Force rates are statistically significantly different from each other (10ms). The grey zone indicates the time during which tactile information was too close to the motor reaction to contribute to detecting the friction condition. **B|** Heat maps of the t-statistics of the differences in strains between catch and normal trials. The t-statistic is set to zero when the difference is not statistically significant and the corresponding p-value is smaller than 0.05. The red color indicates more strains in the case of the catch trial and the blue color more deformations in the case of the normal trial. The first line compares normal high friction trials to catch-low friction trials and the second line compares catch-high friction trials and normal low friction trials. **C|** Average value of the strain rate norm at −150ms normalized w.r.t. the value at the peak of LF, in the different zones of the fingertip indicated in Fig 5B. The brackets show the 95% confidence interval of the mean. The bars delimited with hashed lines indicate the catch trials. One point corresponds to one individual participant.

In Fig 5A, we first show the catch-low and catch-high Δ GF traces zoomed in a 600ms time window centered on the LF peak, which appeared to be the instant of inflection of the curves. The time derivatives of those curves are also shown below in Fig 5A. Note that in this figure, in contrast to Fig 4, we excluded the data from two participants having consistently poor image quality and we also only included the trials in which the images reached high-quality standards (84% of trials among the remaining participants, see *Methods* for more details). As observed in Fig 5A, the two Δ GF traces followed very similar trends just after contact and started to diverge around the LF peak. We found that the Δ GF rates start to diverge significantly 10ms after the time of peak LF (or 370±76ms after initial contact or 176±21ms after liftoff), suggesting that online corrections to friction arise around this moment. Note that this is also approximately the time when the Δ GF rates become statistically significantly positive, or negative, for the “catch-low” trials, or the “catch-high” trials, both 20ms after the LF peak.

The different responses of participants to catch-low and catch-high trials suggest that a sensory signal informative about the frictional properties of the material originated prior to this time. We further needed to take conduction delays into account, which for tactile-motor responses are close to 90ms on average. Accordingly, we drew a grey box that encompasses the 90ms before the instant of significant difference to indicate the minimal interval of time needed for the sensory feedback to reach the central nervous system and trigger a motor response (Pruszynski and Johansson, 2014; Pruszynski et al., 2016; Crevecoeur et al., 2017). Thus, to trigger a motor reaction observable just after the LF peak, a sensory signal must be readily observed before the beginning of this grey box (i.e. 80ms before LF peak).

Accordingly, we evaluated the skin strain associated to partial slip during a period encompassing 300ms before the LF peak until 100ms after (see Supp Fig 4). In the same manner as for the GF, we report here the contrast between the catch and the normal trial, looking for a difference that might have triggered a corrective behavior. In Fig 5B, we report the differences in strain rate norm between the different conditions (catch-low, top row and catch-high bottom row), as expressed in the form of a t-value, obtained from paired t-tests performed for each pixel over the whole fingertip contact area (see Methods). Red zones show the parts of the fingertip where strain rates were significantly higher for the catch trials, i.e. where a significant excess of strain rate was observed, whereas blue zones show parts of the contact having a significantly lower strain rate for catch trials. Green zones depict insignificant differences (t(14) < 2.145, alpha = 0.05). Strains were first normalized according to their average value at the time of peak LF (see *methods* for more details and Supp Fig 3 for normalization values). This normalization procedure was performed because subjects showed markedly different average levels of strains.

For “catch-low” trials shown in the top row of Fig 5B, a zone of excess strain rate norm in the distal part of the center of the fingertip starts appearing 250ms before the maximum of LF and is consistently observed until the LF peaks. This excess disappears progressively when participants start to adapt their GF to the level of friction and when the general level of strain rate becomes small. Conversely, some smaller zones in the periphery show lower levels of strain rates. Thus, surprising the participant with a material with a friction lower than for the previous trials generates a consistently higher level of strain rates deeper inside the contact areas while leaving the periphery less strained at some places. This striking observation is valid over the span of several hundred milliseconds and can thus be taken into account by participants when adjusting their GF.

For the “catch-high” trials shown in the bottom row of Fig 5B, we do not observe any large zone of difference in strain rate norm. Some patches in the more central part tend to show lower levels of strains rate norm. The sensory signal is less contrasted in the case of “catch-high” trials suggesting that the GF decrease might be an automatic slow decrease following a light GF excess applied because of a new block. A typical “catch-high” and “catch-low” trials are provided in the supplementary video. These show that the strain rate reaches higher level deeper in the surface of contact in the case of “catch-low” trials.

The findings are quantified in fig 5C where we arbitrarily defined different zones in the fingerpad, using two ellipses with the smaller one having radii 2/3 as large as the larger one, and quantified the average level of strain rates for each participant at 150ms before the peak of LF and thus approximately 160ms before the first signs of adaptation to the new friction condition. For the exterior part of the ellipse, although the level of strain rate is the highest, there is no statistically significant difference between the two conditions (paired t-test, diff=−0.36±1.4, t(14)=−1.02, p=0.3257). For the central proximal half part of the ellipse, the level of deformation is lower and there is still no statistically significant difference of level of strains between the two conditions (paired t-test, diff=−0.2±0.66, t(14)=−1.12, p=0.251). It is only for the central distal half part of the ellipse that we observed that the amount of strain rate norm was almost twice as high for the low friction catch trials as for the normal high friction trials. This difference is large, consistent among participants and statistically significant (paired t-test, diff=-0.69±0.8, t(14)=−3.37, p=0.0046). We performed the same analysis for the catch-high trials, but observed no statistically significant differences, as can be seen in the right panel of Fig 5C (Exterior ellipse, paired t-test, diff=0.26±0.62, t(14)=1.59, p=0.1343. Interior proximal, paired t-test, diff=0.065±0.4, t(14)=0.6, p=0.5609. Interior distal, paired t-test, diff=0.0705±0.35, t(14)=0.78, p=0.4497).

As the observed deformation contrasts take place before the grey zone defined earlier, they can contribute to the information used by the participants to adapt their GF to the condition of friction.

In summary, we observed the onset of a motor response resulting from the friction change approximately at the moment of the LF peak and this online GF correction was consistent with a sensory signal resulting from an increase in the deformations closer to the central parts for the contact area happening over 100ms before the motor response. This subtle but essential sensory signal, therefore, explains GF adaptation to changes in friction.

## Discussion

This is, to our knowledge, the first study that quantifies how fast humans can adjust their GF to a change in friction in the case of flat transparent surfaces. We demonstrate that this adjustment can be based on a local strain pattern that takes place in the contact area with the manipulated object and signals an insecure grip. Specifically, we show that when confronted with lower friction than expected, skin strains advance deeper and faster in the contact area. The differences in skin strains with respect to a normal trial are already significant very early after the initial increase of the load force, and more than 100ms before the motor response, which is a reasonable delay to explain it (Macefield and Johansson, 2003; Crevecoeur et al., 2017). These differences are present over a period of time of several hundred of milliseconds and can thus constitute a warning signal that allows the central nervous system to adjust the GF to the friction condition.

We observed that the levels of skin strains varied greatly from participant to participant, as can be seen by the strain rates normalization values in Supp Fig 3. We have shown similar levels of variation of skin strains in a previous study where participants had to perform oscillations in a precision grip (Delhaye et al., 2021b). We can also observe in that study that a larger amount of skin strains is linked to a lower stick ratio and that the stick ratio also varied greatly between participants. The stick ratio was also measured in a study where participants had to perform a grip-lifting task (Tada et al., 2002). The authors hypothesized that humans control the level of GF to maintain a constant amount of partial slip (~40% of the contact area) when lifting objects of known friction and weight, but they mentioned in their paper that the validness of this hypothesis has to be verified, as only three participants were tested. It is worth noting that humans perceive slippage at very different levels of partial slip in a passive setting (Barrea et al., 2018). The variability in the levels of strains and stick ratio during manipulation and the variability in partial slip between participants seem to point towards strategies of manipulation that vary from person to person. This requires further inquiring, by performing experiments with tasks of different nature, with varying friction and weight of the manipulated object. A characteristic of our results is the asymmetry of the participants’ behavior between “catch-high” and “catch-low” trials (Fig 4). Although the motor reaction seemed to be triggered at the same time in both conditions (Fig 5A), it took more time to reach a level of GF that was adapted to friction in the catch-high condition. It is worth noting that a short reaction time in the catch-low condition is critical: it is urgent to increase GF since not correcting it could lead to a dramatic slip and drop of the object. In contrast, the excessive GF in the catch-high trials only results in a temporary slight excess of energy expenditure, which does not require to act quickly. The difference in urgency to adapt GF between conditions is observed in the differences of strains (Fig 5 B-C), where a surprisingly low level of friction caused significant differences in strains whereas a surprisingly high level of friction did not.

Although we characterized the friction between the fingers and the manipulated object by a constant scalar value per participant-condition pair (Fig 2B-C), this is clearly a gross approximation of a very complex phenomenon. The tribology of the skin is complex and varies significantly at different temporal and spatial scales (Pataky et al., 2005; Adams et al., 2013; Van Kuilenburg et al., 2015). One aspect of this complexity is related to the complex geometry and mechanics of the finger: the fingertips are composed of several layers, from the bone in the interior of the fingertip to the epidermis in the exterior. The epidermis of the glabrous skin is characterized by the presence of ridges and furrows that form the fingerprints and present a complex topography (Choi et al., 2021). This complex geometry and mechanics is likely to impact the friction on a trial to trial basis, depending on how the finger contacts the object. Another aspect of the complex tribology of the skin is that it evolves over time: Sweat pores are present at the surface of the skin and produces humidity at the interface that heavily impacts the level of friction when gripping a material, which changes over the course of several seconds (André et al., 2010; Yum et al., 2020). The occlusion phenomenon defines humidity evolution and plasticization of the skin when in close contact with the material for a few seconds (Adams et al., 2013). The level of friction thus varies during a single trial as moisture varies due to occlusion. In our experiment, the moisture level is however probably partially maintained when trials are close enough from one another. Thus, in the context of this study, the differences in friction measured between materials, which could vary widely between participants, are to be considered as approximations of their actual values, which can vary during manipulation. However, it is clear that the difference in friction between materials was consistent and sufficient to generate coherent adaptation of GF during the overall experiment.

Several of our senses are used when performing a task as difficult as fine object manipulation. Our sense of touch in particular is very important. It sends critical information to the central nervous system to adjust GFs to various parameters of the manipulation such as friction. We have shown that when gripping and lifting small objects, the skin strains depended on the level of friction at the interface of the contact. The localized differences in skin strains between conditions during the loading phase were consistent with the timing of the first signs of adaptation of the GF and were sufficient to explain them. We have quantified and described those skin strains and measured the GF modulation, which started less than 500ms after the initial contact.

## Methods

### Participants

Eighteen volunteers (5 females; ages 20-65) participated in the experiment. All of them provided written informed consent to the procedures and the study was approved by the ethics committee at the host institution (UCLouvain, Brussels, Belgium).

### Apparatus

At rest, the device was standing on a table with a hole to allow the passage of the cables coming from the bottom of the device. Its weight (540g) was partially compensated by a counterweight (320g) attached to a system of pulleys. The device is described at length in a recent publication(Delhaye et al., 2021b). Succinctly, forces were measured under each fingertip using two six-axis force and torque sensors (ATI Mini27 Ti, ATI-IA, Apex, NC, USA). From those measurements, the GF and LF were computed, as described in *Data Analysis*. The position was measured using an optical distance sensor (DT20-P224B, SICK Sensor Intelligence). The position and forces were sampled at 200 Hertz with a NI-DAQ card (NI6225, National Instruments).

A custom optical system allowed to image the index fingerprints in contact with the glass (Fig 1F). Because of constraints in the design of the manipulandum, it was only possible to monitor one side, as the light is emitted by an array from the side where the thumb is, blocking its observation. This system is based on the principle of Frustrated Total Internal Reflection and enables a high contrast between the point in contact with the glass and those that are not. Images are recorded at 100 fps with a camera (GO-5000M-PMCL, JAI, monochrome, 2560 × 2048 full pixel resolution). Image size is 1696 × 1248 pixels with a resolution of 4096 pixels/mm^2^, which corresponds to an area of 26.5 × 19.5mm.

Two kinds of glass plates were used to generate different levels of friction. The first set of plates are simple transparent optically flat plates of glass. They are referred to as “high friction”. A process called “glass frosting” was used to alter friction in the second set of plates. In brief, a chemical process was used to imprint a nanoscale pattern on the surface of the glass. With the right set of parameters (height and roughness), this decreased the real area of contact between the finger and the plate and thus the coefficient of friction (Derler et al., 2009; Skedung et al., 2010, 2011; Adams et al., 2013; Wiertlewski et al., 2016; Inamoto and Tomeno, 2019). This nano-structured glass was referred to as the “low-friction” surface. The transparent plates are indistinguishable to the naked eye.

### Experimental procedures

Participants stood in front of a table on which the device was positioned. After an auditory cue, they were instructed to grip and lift the device to a height of about 20cm within 0.8s, and then hold it still for 1.5s. They then performed three fast point-to-point movements (0.8s) with pauses (1.5s) in-between. Auditory cues were used to pace each movement. We observed that participants’ movements were slightly slower than what was instructed, resulting in slightly longer movements and shorter static phases. The participants were requested (and often reminded during the experiment) to use a minimal amount of GF. Indeed, we observed in a preliminary study that participants tended to use an excessive amount of GF naturally, probably because this device composed of a camera and sensors seems fragile and looks heavy. The glass plates were cleaned with alcohol after each trial. This served the purpose of getting images as clean as possible. Also, this procedure removed sweat that could alter the topography of the glass plates at a microscopic level and thus the level of friction. After each block of five trials, participants were instructed to take a break on the other side of the room, from a location where they could not see the experimenter manipulating the device. During that break, the experimenter interchanged the plates such that the friction was changed from high to low or from low to high. This procedure was quick and took a maximum of 2 minutes. Half of the participants started with the high friction condition and the other half with the low friction. This caused no difference in their adaptation to friction, as measured by the difference in GF between conditions during the first static phase of normal trials (t-test, t(15)=0.36,p=0.72). The coefficient of friction was measured for each material at the end of the experiment (see *Friction Measurement*). In total, the experiment lasted between one and a half and two hours for each participant.

#### Friction measurements

We measured the coefficient of friction between the participants’ fingers and both materials at the end of the experiment using the method described in (Barrea et al., 2016). Briefly, participants were instructed to rub their index finger and thumb on the glass plates for three periods of fifteen seconds at different levels of normal forces. The approximate range of normal forces for each period was 0 to 2.5N for the first, 2.5 to 6N for the second, and 6 to 10N for the third. The moment of slippage was detected by finding the maximum of the ratio of tangential force over normal force at the start of each rubbing motion. This ratio was measured and was our estimation for the static coefficient of friction corresponding to the normal force applied at that moment. The data were obtained for both fingers of all participants (see Supp Fig 1–2) and fit with a negative power law (André et al., 2009). From the fits, we computed a single coefficient of friction value for each participant and each material for the 1-5N range that corresponded to the approximate range that the participants used for manipulation. The friction was averaged across both fingers. The measurement of the friction was always performed first on the material with which the participant finished the experiment.

### Data analysis

#### Forces and position

data were filtered with a fourth-order low-pass Butterworth filter with a cut-off frequency of 40Hz. The GF is defined as the mean of the norm of the forces normal to the surface of the object exerted by each finger. The LF is defined as the norm of the sum of the forces applied tangentially to both surfaces by each finger. When comparing dynamics or kinematics between participants as in Fig 3, 4 and 5, an average curve was first computed for each participant by first synchronizing all trials on the maximum of LF for each separate movement. Then, statistics such as the inter-subject mean or standard error of the mean were computed based on these average curves.

#### GF normalization

As participants typically used different levels of GF, a normalization procedure was used when comparing the GF signals across conditions and participants. A GF normalization value was obtained for each participant by taking the mean of GF during the first static phases of all normal trials (except the first two blocks that were excluded because of the learning period). They are shown in Supp Fig 3.

#### Image processing

A previously described image processing pipeline was used to evaluate the skin strains from the raw images (see Delhaye et al, 2014, 2016). This pipeline is only summarized here. First, a custom-made machine learning algorithm was used to detect the area of contact between the finger and the glass plate for each image. This algorithm was trained for each participant separately with manually detected areas of contact for randomly selected images. Then, feature points were selected automatically from several frames of the sequences of images (at the beginning, during, and at the end of each movement). Their position was tracked forward and backward in time from frame to frame using an algorithm of optical flow (Lucas and Kanade, 1981; Shi and Tomasi, 1994). A Delaunay triangulation was then computed and the evolution of the shape of the triangles allowed to measure the local strain rate along three dimensions (vertical, horizontal, and shear strain rate). The norm of the strain rate was calculated as follows:

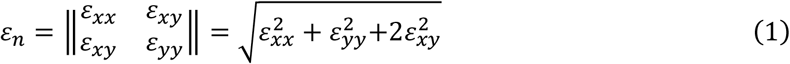

where *ε*_*xx*_, *ε*_*yy*_ and *ε*_*xy*_ are the horizontal, vertical, and shear strains rates components respectively. This gave a quantitative measure of how the different parts of the fingertips were being deformed, irrespective of the type and directions of these deformations. In this study, we were mostly interested in comparing the amount of strain rate according to the condition of friction and the adaptation of GF rather than the specific description of these deformations(Delhaye et al., 2021b).

As different participants showed markedly different levels of strains, we normalized the strain rate norm of each participant by the mean value of the strain rate norm across the entire area of contact and trials at the time of maximum LF. The normalization values of the strain rate norm are given together with the normalization value of the GF for each participant in Supp Fig 3.

The first images of the contact were difficult to interpret. Indeed, the fingertip skin can be rough and stiff on a small scale, depending on the moisture content of the individual’s skin. When it enters in contact with a stiff surface such as glass, the initial real area of contact is low. However, during the first tens of milliseconds of the contact, moisture secreted by the sweat pores hydrates the skin, rendering it softer and elasticizing it. The skin then enters in closer contact with the surface and the real area of contact increases (Pasumarty et al., 2011; Bochereau et al., 2017; Dzidek et al., 2017). As a rapidly changing real contact area not associated with skin deformations was problematic for the interpretation of the results of our image-processing pipeline, we decided to discard the images directly following the time of contact between the skin and the surface from our analysis. We used the first image of the loading phase, defined as the moment when a participant starts applying tangential force to lift the object after the pre-loading phase (Westling and Johansson, 1984) as the first image in our image processing pipeline. This guaranteed that the apparition of moisture would only play a negligible role in our measurement of strains and that the strains caused by the vertical lifting of the object would be included in our analysis.

#### Strain rate summary across participants

To get a summary of strain rates across trials and participants, we first had to project the values from the triangulation used to compute the strains to a standardized structured grid common for all participants. To that end, we first created a reference ellipse with a major to minor axes radii ratio of 3/2. A gridded mesh of size 91×61 was attached to this reference ellipse. Then a least-square procedure was used to fit ellipses on the coordinates of the fingerprint contact area contour for all individual images (Fitzgibbon et al., 1999). Finally, we computed the projection (offset, scaling, and rotation) from the ellipse obtained from the image at the instant of the LF peak to the reference ellipse for each trial and movement. This projection, obtained for each trial and movement, was applied to the center of each triangle for all images to obtain strains on the reference ellipse. The maximum of LF was chosen to compute the projection because this timestamp is used to synchronize the trials. At that instant, the area of contact between the finger and the glass has already plateaued. After having applied this projection, the strain rate norm was averaged for each participant and each condition (i.e. catch/normal and low/high friction). An example of such data for a typical participant is presented in Supp Fig 4. Subsequent statistics on strains across participants were performed on the projected data.

#### Inspection and sorting of trials

As mentioned in *Results,* the first two blocks were considered to be “training blocks” as the participants’ GFs decreased significantly during those for all participants and were thus excluded from the data analysis (Fig 1C). The third block was the first included in our analysis because it was the first for which (1) the grip forces were on average smaller than in the next one in the high friction condition and (2) the grip forces were larger than in the next one in the low friction condition. For some trials, participants placed their index finger outside of the field of the camera or displaced it outside of the field during the trial due to slipping or rolling. Some trials were therefore not included in the analysis. Only the parts of the trials where the finger got out of the field of the camera were removed. After a close inspection of each trial, 99 out of 720 were at least partly removed because the images were unusable during some part of the trial. Those trials were still used for the kinematics and dynamics analysis of Fig 2–4. Also, 15 participants were used for the image and forces analyses of Fig 5 because two participants had very dry skin and the image quality was insufficient to obtain reliable strain data.

#### Full slip trials

To get a different look at our data, we counted the proportion of trials for which a full slip occurred during the first movement, which yielded the following results: a full slip occurred in 11.33% of the high friction normal trials, 12.75% of the low friction normal trials, 11.77% of high friction catch trials, and 31.03% of low friction catch trials. A full slip is said to occur when all the feature points whose positions are tracked from frame to frame are measured as moving with respect to the glass. This result is consistent with the differences in strain that are shown in Fig 5 B and C. That is, in some cases, the strain wave reached the central point of the contact. Note that even if full slip occurred for some trials, the extent of slippage was small and was quickly stopped by a corrective GF (slipping distance of the central part of the contact area of 0.19±0.25mm, mean±std, for trials in which the full slip was reached). The full slip trials were included in the strain analysis like all other trials.

### Statistical analysis

All statistical analyses were performed with MATLAB, using the functions *corr* (for Pearson correlation), *ttest* (for paired t-tests), and *tinv* (inverse of Student’s T cumulative distribution function). The test performed, the number of degrees of freedom, and the T-statistics are always mentioned with the p-value. The 95% confidence interval of the means were computed using one mean value per participant and using the following formula:

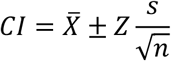

where CI is the confidence interval, 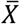 is the average across participants, Z is the inverse of Student’s T cumulative distribution function (e.g. Z=2.12 for 17 participants), *s* is the standard deviation and n is the number of participants.

## Acknowledgments

The authors thank Allan Barrea, François Wielant and Arsalis for their help in the development of the device. This work was supported by a grant from the European Space Agency, Prodex (BELSPO, Belgian Federal Government). BPD is a postdoctoral researcher of the Fonds de la Recherche Scientifique – FNRS (Belgium).

---

**Supp. Figure 1:**
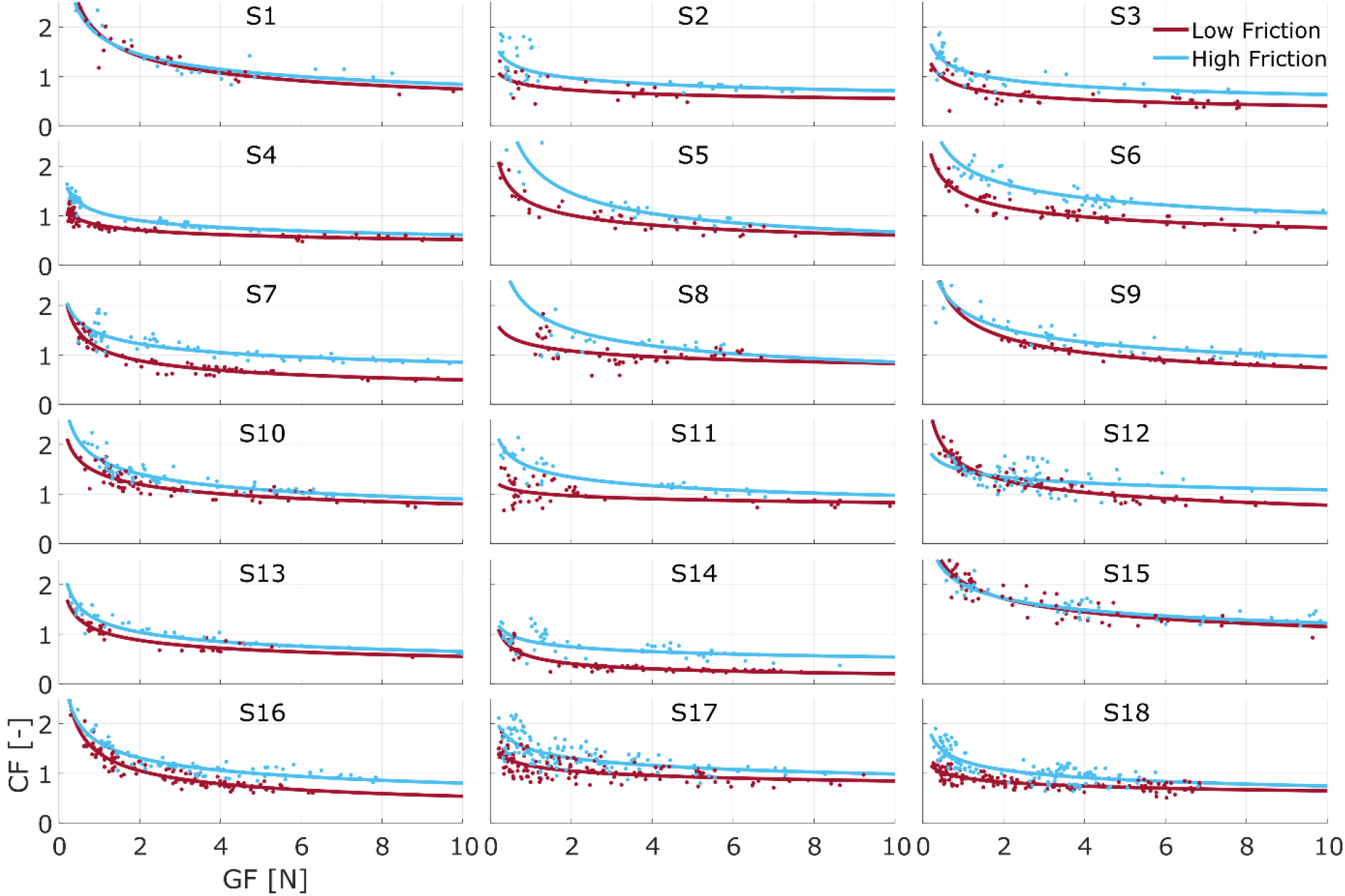
Coefficient of friction of the index finger for all participants.

**Supp. Figure 2:**
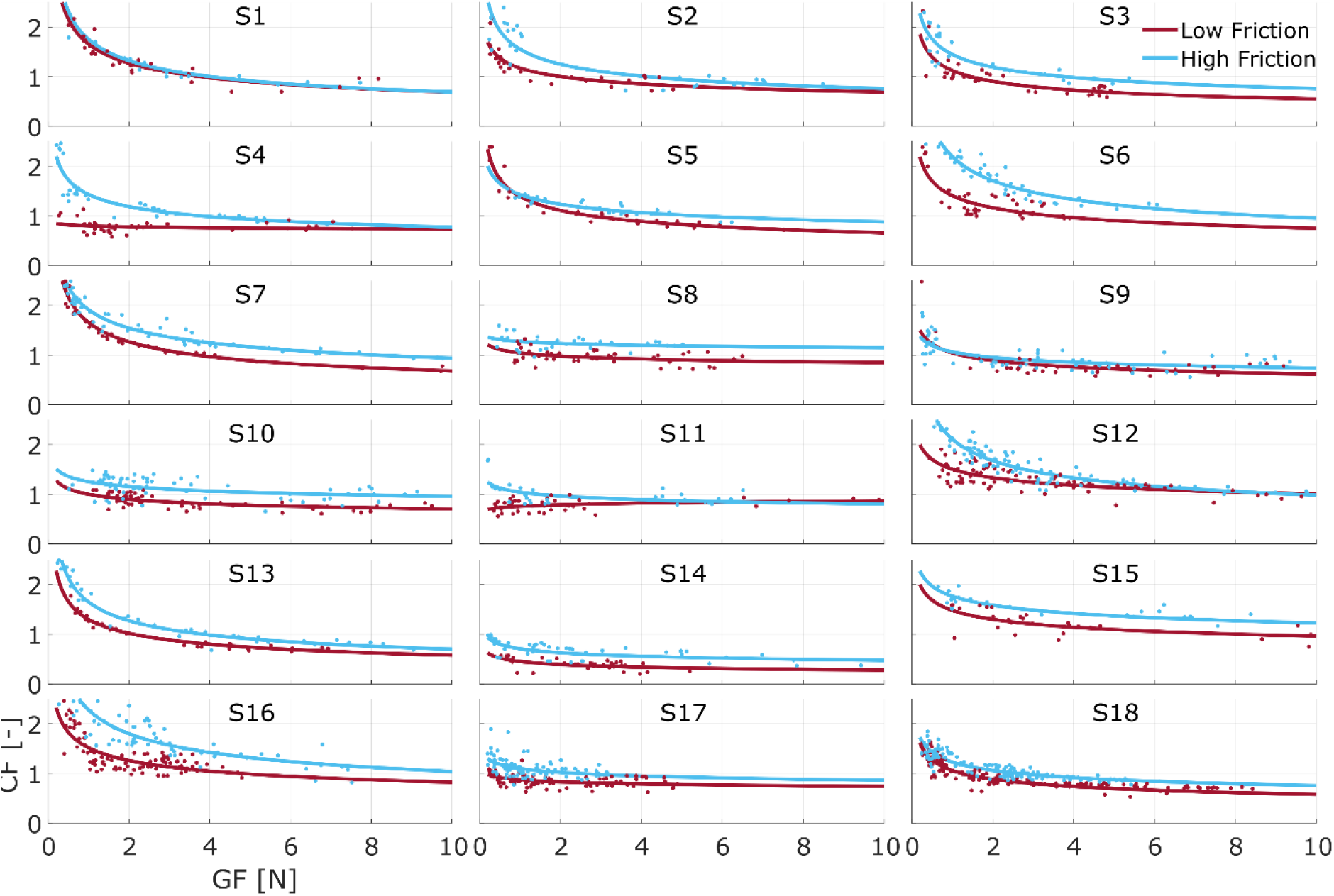
Coefficient of friction of the thumb for all participants.

**Supp. Figure 3:**
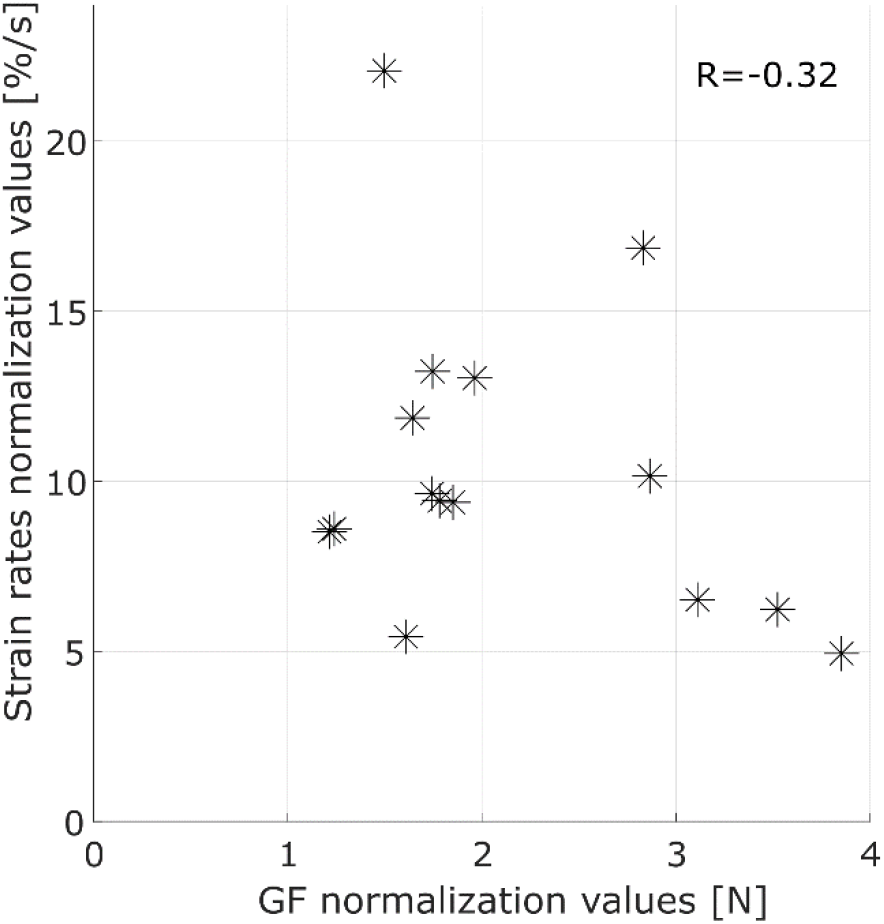
Normalization values for the GF and strains.

**Supp. Figure 4:**
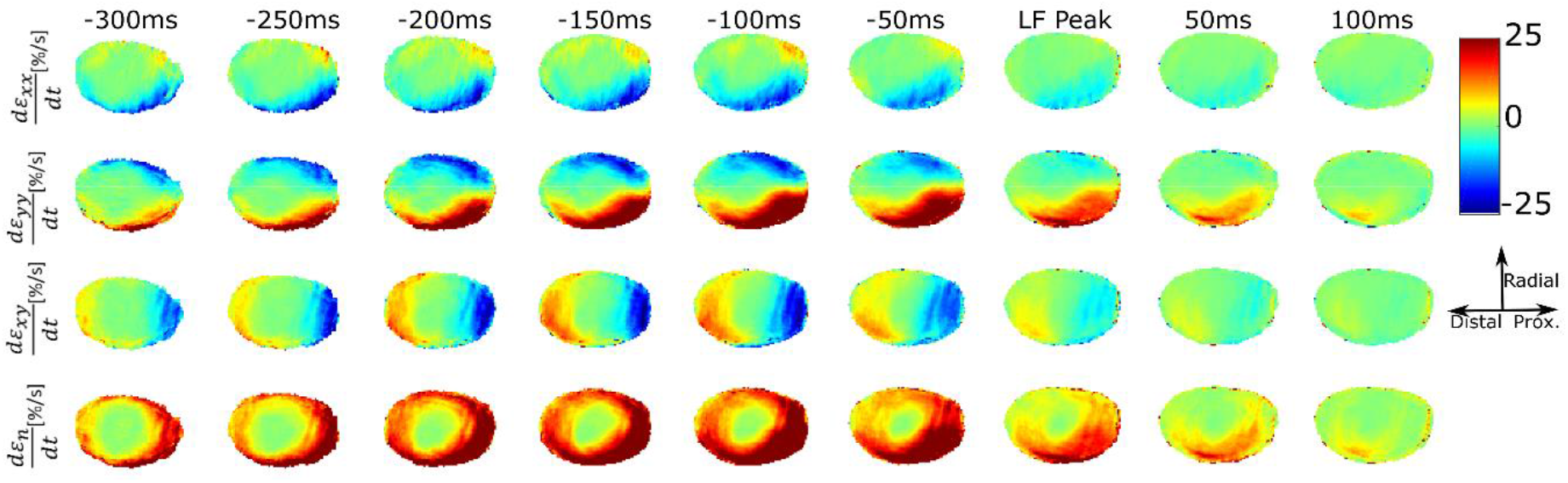
Averaged strain rate data on the standard grid during the first movement for a typical participant. Compression is shown in red and dilation in blue. The first row shows the horizontal strain rate, the second shows the vertical strain rate, the third shows the shear strain rate and the fourth shows the strain rate norm. The average was computed over all normal high friction trials (n=15) for a typical participant.

